# Controlling metal-carbonate phase, form, and function through *de novo* protein design

**DOI:** 10.64898/2026.06.10.730916

**Authors:** Paul S. Kwon, Xinqi Li, Le Tracy Yu, Harley Pyles, Todd H. Lewis, Connor Weidle, Andrew J. Borst, Catherine C. Bodinger, Alex Kang, Hannah Nguyen, Johnny Mendoza, Kenneth D. Carr, Brian Coventry, Yang Hsia, Zsombor Molnár, Dongsheng Li, Bo Zhang, Brandi M. Cossairt, Shuai Zhang, Asim K. Bera, James De Yoreo, David Baker

**Author notes:** These authors contributed equally: Paul S. Kwon, Xinqi Li.

## Abstract

Biomineralization enables living systems to construct hybrid materials by controlling the location, orientation, and polymorph of inorganic crystals with proteins and other biomolecules. Despite decades of study, the molecular principles underlying these processes remain difficult to harness in engineered materials, in part because native biomineralization proteins are often intrinsically disordered, heterogeneous, or insoluble. Here we show that de novo designed protein interfaces can be assembled into reconfigurable two-dimensional arrays which template calcite nanocrystals. By fine-tuning RFdiffusion2 on repeat protein scaffolds, we further enable the design of protein architectures which selectively form aragonite, a metastable polymorph of calcium carbonate, in nucleation conditions that otherwise result in a mixture of phases. Extending beyond inorganics found in biological systems, we show that lattice-matched protein designs template cobalt carbonate formation: a flat helical repeat protein interface promotes unconfined growth, whereas soluble D3 cage assemblies yield more homogenous cobalt carbonate nanocrystals confined to the interior of the cage. These protein-cage cobalt carbonate hybrid materials function as electrocatalysts for alkaline water splitting. Our results demonstrate the potential of deep learning-based methods to unlock the structural and functional activity of protein-mineral composites.

## Introduction

Biomineralization enables living organisms to fabricate functional materials with exquisite precision, exemplified by shells^1^, bones^2^, teeth^3^, and magnetosomes^4^. In mollusk shells, calcium carbonate (CaCO_3_) deposition is controlled by organic matrices of chitin and proteins such as Pif^5^, yielding composites with hierarchical organization^6^, and polymorph selectivity^7^ that results in a fracture toughness^6^ greatly exceeding the mineral alone. Understanding the design rules underlying proteins that orchestrate biomineralization could enable the development of synthetic biomimetic materials with applications in sustainability^8^, biotechnology^9^, medicine^10^, and energy^11^. However, key features of biomineralization remain unrealized with designed proteins^12^, including stabilization of metastable polymorphs, intentional targeting of non-natural crystal faces, templating across protein arrays, and confinement within enclosed volumes. Achieving synthetic control over these features would enable extension to non-natural materials and functions, such as electrocatalytic protein-mineral composites, that require mineral-directing interactions to be encoded not only in individual protein monomers, but also across higher-order oligomeric architectures.

Native biomineralization proteins are often intrinsically disordered, heterogeneous, or insoluble, posing substantial challenges for structural analysis and structure-based engineering^13^. Previously, we showed that these limitations could be overcome by creating an epitaxial match between structurally defined *de novo* proteins^14^ and crystal lattices^12^ to encode a programmable mineral-binding interface^15^ that controls mineral nucleation, growth and facet selection. Using designed helical repeat proteins (DHRs) we exploited this lattice-matching approach originally inspired by native ice-binding proteins^16^ to create templates that direct nucleation of calcite^15^, hydroxyapitite^17^, zinc oxide^18^, iron oxide^18^, and ice^19^. Despite these advances, it remains an unsolved challenge to design protein architectures that stabilize metastable mineral polymorphs, organize mineral growth on reconfigurable two-dimensional lattices, and confine crystallization within shape-defining enclosures.

Here we set out to overcome these limitations by designing protein systems that promote the formation of metastable polymorphs, mineralize over arrays spanning tens of nanometers, and template mineralization within closed containers that define the shape of the forming nanoparticle. We focused on metal carbonates as particularly compelling targets because they are central both to the biogeochemical carbon cycle^20^ and to broader materials applications^21^.

### Machine-learning (ML) design methods for inorganic mineral templating

We began by developing ML methods for designing proteins for inorganic mineral templating, leveraging the recent advances in generative protein design using RFdiffusion, and in atomic motif scaffolding^22^. We started from RFdiffusion2 (RFD)^22^, which enables direct generation of protein structures which precisely scaffold pre-specified atomic motifs interacting with non-protein atoms. RFD2 was developed for enzyme design, where the non-protein atoms are the reaction substrates and transition states, and the atomic motifs are the protein catalytic machinery. In the design of mineralization, the geometry is quite different; the non-protein atoms are in a coherent crystal array which extends over a length scale longer than a protein molecule (rather than a reaction substrate which typically only interacts with a small part of the protein) and the atomic motifs are regularly arranged over an extended region of a crystal surface, rather than confined to a localized active site (**Fig. 1a**). Furthermore, for enzyme design the goal is to generate a globular protein with an active site pocket hosting non-protein atoms, whereas in design of mineralization, proteins with repeating substructures that match the periodicity of the inorganic crystal lattice are desirable because the size of the mineralizing interface can be readily increased or decreased by adding or removing repeat units, and because an optimal solution to the crystal templating challenge need only be found in one local region and can then be readily propagated. To optimize RFD2 for non-protein ion templating, we incorporated conditioning on secondary structure and adjacency matrices, and included a large set of repeat proteins derived from the PDB^23^ and AlphaFold (AF) database^24^ for training (composition of training set 25% repeat proteins, 75% the overall PDB). Once trained, RFD2-repeat protein (RFD2-RP) generated novel protein-mineral interfaces in the form of repeat proteins composed entirely of α helices, β sheets, or mixed secondary structure (**Fig. S1**). To bias designs toward fully helical architectures, 70% of each sequence was specified as α helices, while the remaining positions were left masked. This strategy produced predominantly helical designs, avoiding β-sheet–rich architectures, which can be prone to aggregation (**Fig. 1b**).

**Figure 1:**
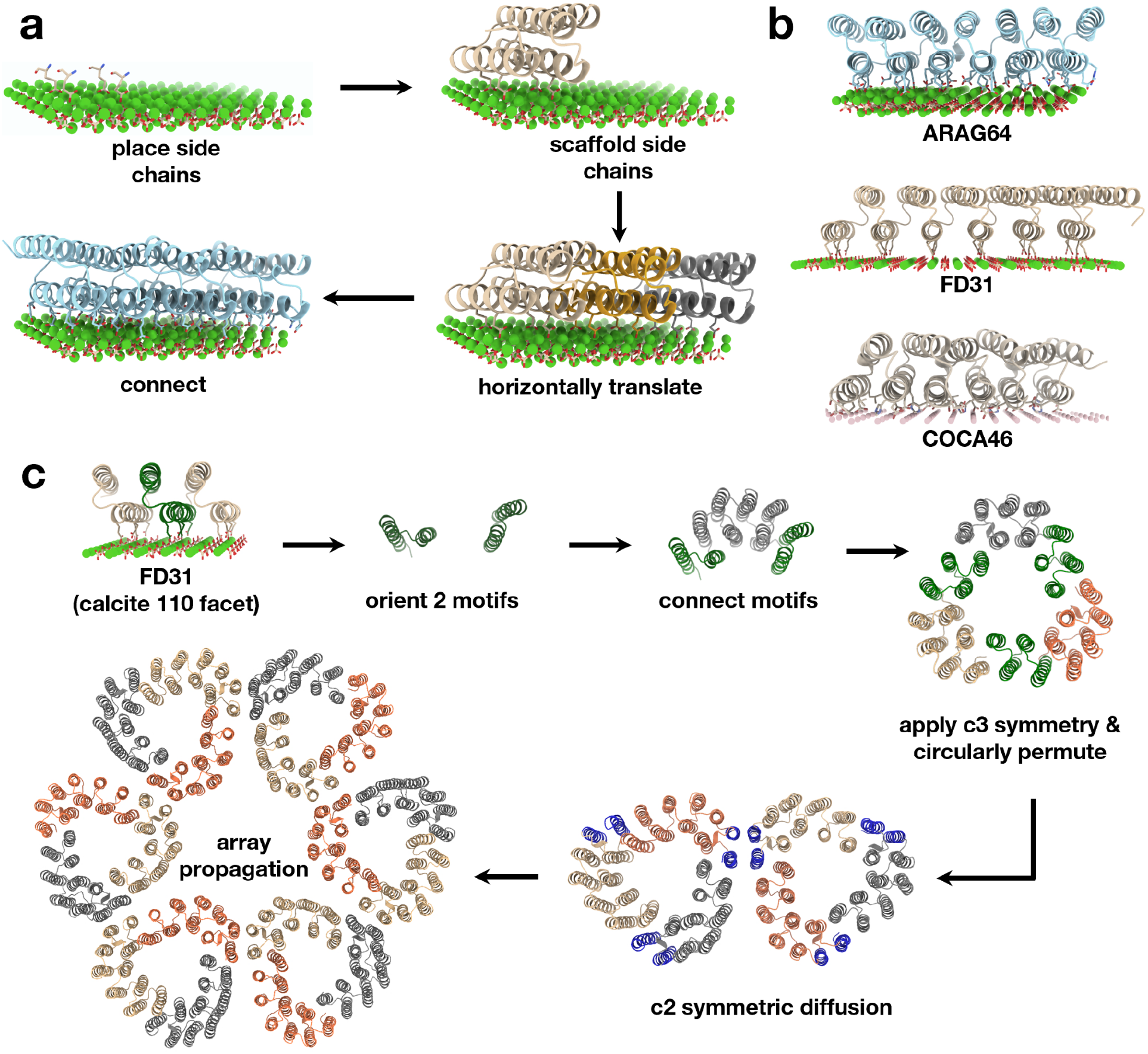
Design of protein–mineral interfaces and higher-order mineral-templating assemblies. **a**, Computational design strategy for generating extended protein–mineral interfaces against the aragonite {110} facet. Repeat-protein scaffolds were generated using a fine-tuned RFdiffusion2 model that was trained on repeat proteins from the PDB and AlphaFold databases. **b**, Representative designs of de novo protein–mineral interface, ARAG64, FD31 and COCA46, targeting aragonite, calcite and cobalt carbonate, respectively. **c**, Strategy for converting the FD31 calcite-binding motif into an assembly array. FD31 protein–calcite motifs are shown in green and were used to design C3-symmetric triangular building blocks, followed by circular permutation and the design of a C2-symmetric inter-triangle interface, shown in blue, to assemble the array.

### Design and characterization of protein-aragonite interface

To selectively template aragonite, a metastable CaCO_3_ polymorph, we designed proteins complementary to the {110} facet (**Fig. 2a**)^25^. Terminated {110} facets are more prevalent than {010} facets in the pseudohexagonal aragonite habit, and present carbonate groups in a geometry more readily matched by acidic amino-acid side chains on a designed protein surface. We positioned glutamate residues so that the oxygen atoms at the end of each carboxylate group aligned with the oxygen atoms of carbonate groups on the mineral surface, then used tip-atom guided RFD2-RP to generate proteins capable of scaffolding these carboxylate groups at the ends of aspartate or glutamate sidechains. To build larger interfaces, we selected designs capable of propagating along the mineral facet through lattice-matched translations without steric clashes, then linked the propagated copies through another diffusion step (**Fig. 1a**).

**Figure 2:**
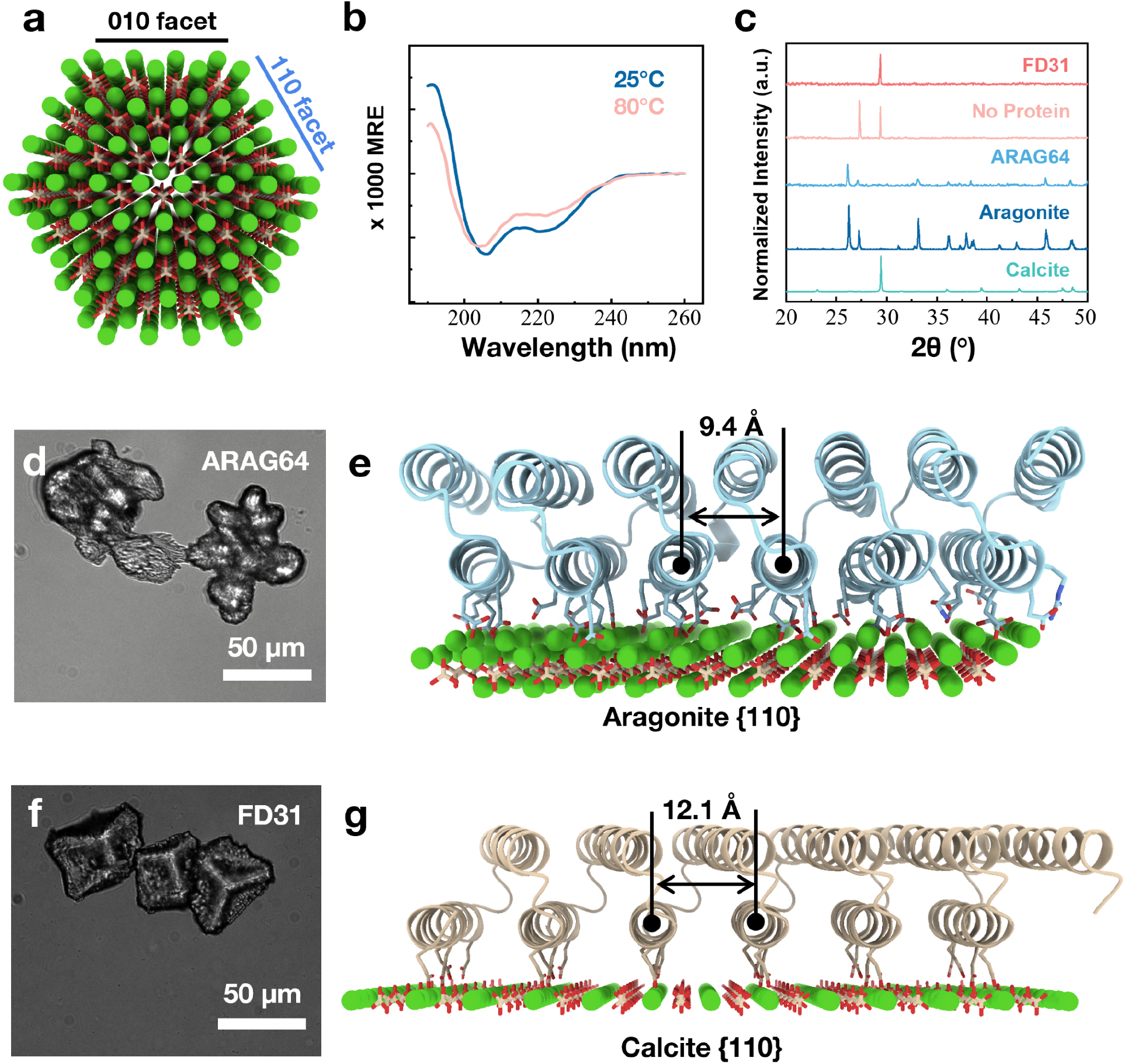
Aragonite and calcite templated by designed proteins. **a**, Atomic model of aragonite viewed along the c-axis, highlighting the {010} (black) and {110} (blue) facets. **b**, CD spectra of ARAG64 protein showing the α-helical structure at both 25°C and 80°C. **c**, Normalized XRD patterns of CaCO_3_ crystal formed in the presence of ARAG64 and FD31 proteins, compared with a no protein control and the references of aragonite and calcite from RRUFF entries of R060195 and R050130, respectively. No-protein controls show a characteristic aragonite peak near 27.2°, assigned to the {021} reflection, and a calcite peak near 29.4°, assigned to the {104} reflection. **d**, Optical microscopy image of aragonite crystals templated by ARAG64 protein. **e**, Computational design model of ARAG64 protein positioned on the aragonite {110} surface, with the designed binding motifs at a distance of 9.4Å for lattice match. **f**, Optical microscopy image of calcite crystals templated by FD31 protein. **g**, Computational design model of FD31 protein positioned on the calcite {110} surface, with designed binding motifs at a 12.1 Å surface-matching spacing.

To represent the aragonite surface during sequence design, we sampled protein sequences in the presence of the Ca^2+^ sublattice on aragonite {110} using LigandMPNN^26^ while keeping the previously designed carboxyl-side-chains fixed. Three designs which AF2^27^ structure predictions were close to the design models were tested in E coli (**Table S1**), two of which were solubly expressed. One design, ARAG64, behaved as a monomer in size-exclusion chromatography (SEC) (**Fig. S2**) with circular dichromism (CD) exhibiting alpha-helical folds both at 25°C and 80°C (**Fig. 2b**).

We next investigated the effect of ARAG64 on CaCO_3_ mineralization. Following incubation of 2.16 µM protein, 10 mM CaCl_2_, and 10 mM NaHCO_3_ at 80°C and pH 8.2 for 2 hours, X-ray diffraction (XRD) showed that crystals formed with ARAG64 matched an aragonite reference pattern, whereas crystals formed with FD31, a protein previously shown to drive calcite nucleation at low temperature, matched a calcite reference pattern (**Fig. 2c**). Optical microscopy further supported these assignments: ARAG64 produced needle-like aragonite crystals (**Fig. 2d**), consistent with the ARAG64 design model targeting aragonite with a 9.4 Å spacing between the indicated helices (**Fig. 2e**); while FD31 produced rhombohedral calcite crystals (**Fig. 2f**), consistent with the FD31 design model targeting calcite with a corresponding 12.1 Å spacing (**Fig. 2g**). Optical micrographs and XRD patterns of the no-protein control exhibited mixed polymorph formation under these conditions with a mixture of rhombohedral calcite and needle-like aragonite (**Fig. S3**) generating both a characteristic aragonite peak near 27.2°, assigned to the {021} reflection, and a calcite peak near 29.4°, assigned to the {104} reflection (**Fig. 2c**). No crystals were observed, and no diffraction peaks were detected in bovine serum albumin (BSA)-containing solutions. One possible explanation is that BSA, which has a reported melting temperature of 63°C^28^, unfolds at 80 °C and nonspecifically binds or sequesters ions, thereby disrupting the ion preorganization required for mineralization (**Fig. S3-S4**). Together, these results demonstrate that de novo designed proteins can direct polymorph outcome in CaCO_3_ mineralization.

### Templating mineralization over designed 2D arrays

Having shown that designed proteins can control CaCO_3_ polymorph selection, we next sought to direct CaCO_3_ organization over longer length scales by designing 2D protein arrays incorporating a CaCO3 mineralizing interface. To complement the aragonite promoting design in the preceding section, we chose to focus on templating calcite. To construct a triangular scaffold, calcite-nucleating repeat protein FD31^15^ motifs were pre-aligned along specified eigenvectors to direct repeat propagation, then translated and rotated under C3 symmetry to define the target geometry. This pre-placement preserved the symmetry relationship between motifs while allowing RFD2-RP to connect them into continuous monomers (**Fig. 1c**). Because the intervening hydrophobic repeat interface is degenerate, the same packing interaction could be reused between adjacent monomers along each side of the triangle, enabling assembly of the intended C3-symmetric triangular architecture.

Next we designed an additional C2 interface onto the C3-oligomer to form a 2D hexagonal lattice. The triangles were circularly permuted to allow the mineralization motifs in the interior of the triangle to be part of one chain, the permuted triangles were positioned close together and a new C2 symmetric interface was diffused^29^. We designed a p6 hexagonal lattice in which the direction of the local C3 symmetry axes alternate or “flip” between nearest neighbors, correcting for out-of-plane curvature and balancing strain throughout the array. The resulting backbones were subjected to ProteinMPNN^30^ sequence design while fixing key glutamate residues in the calcite interface from FD31. Designs for which monomer predictions with AF2 matched the monomer design model, and hexamer predictions with AF3 were close to two adjacent lattice new cells, were selected for experimental characterization.

Of 96 designs screened experimentally (**Table S1**), one, CALC_HEX24 (**Fig. 3a**), formed well-ordered two-dimensional arrays. Arrays formed in samples containing 3 mg/mL protein and of 500 mM NaCl, and separated into a dense bottom fraction after overnight incubation at room temperature (**Fig. S5**). Assembly was confirmed by negative-stain transmission electron microscopy after ∼457-fold dilution of the bottom phase into a buffer containing the same salt concentration (**Fig. 3b**). Removal of NaCl by dialysis resulted in reversible disassembly of the arrays (**Fig. 3c**) Two-dimensional class averages of the assembled arrays were processed in cryoSPARC^31^ and revealed a hexagonal lattice composed of repeating triangular units (**Fig. 3d**). The disassembly of the arrays at low salt (likely due to electrostatic repulsion between the highly charged monomers) enables convenient soluble expression and purification of the single-component array, in contrast to typical one-component arrays that accumulate in insoluble inclusion bodies and require denaturation and refolding^32^. This salt-dependent assembly combines the practical advantages of single-component systems with key benefits of two-component arrays^33^, in which assembly is induced by mixing two individually soluble subunits.

**Figure 3.**
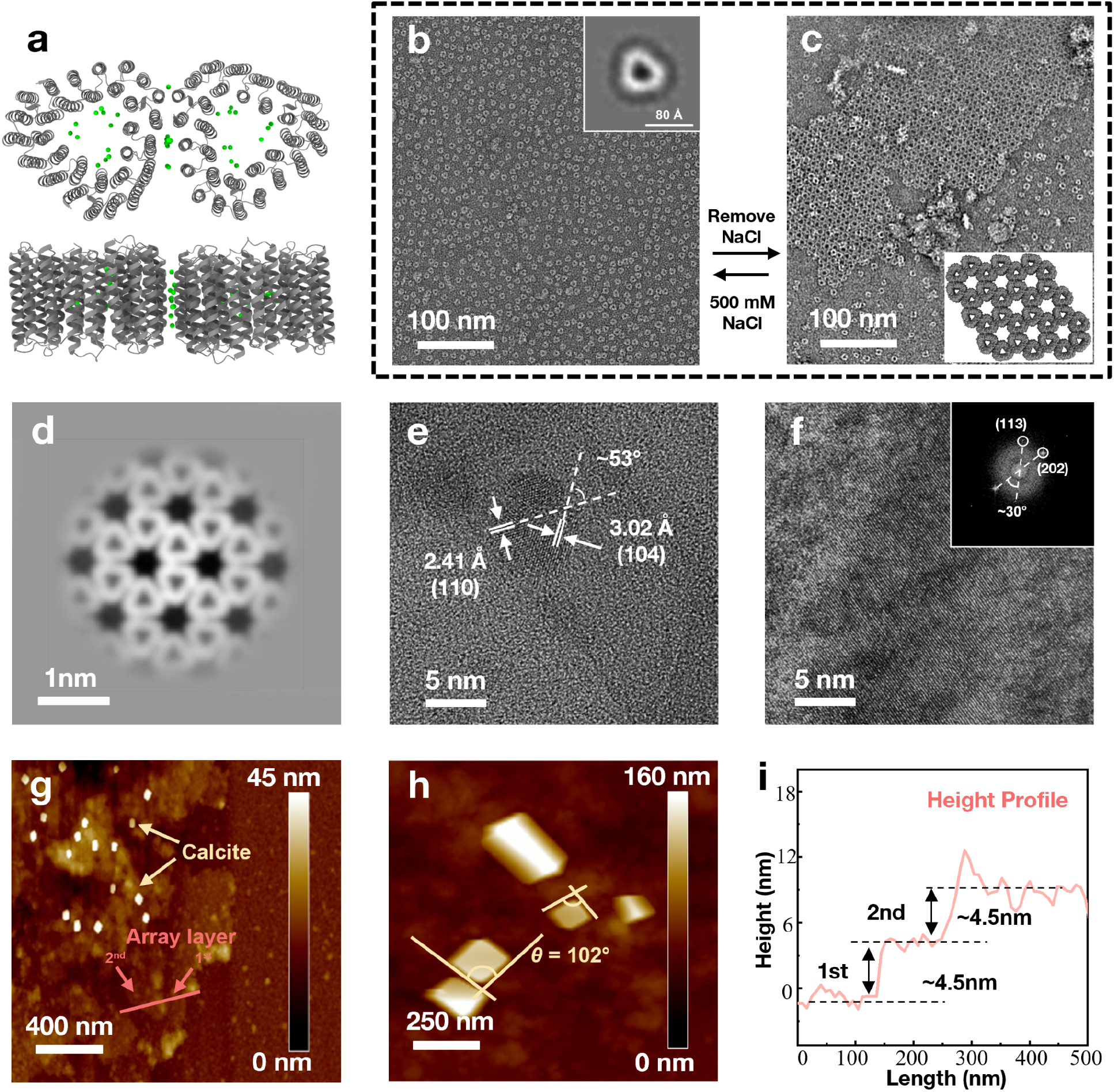
Calcite biomineralization templated by reconfigurable CALC_HEX24 arrays. **a**, Axial and lateral views of the AlphaFold 3 model of CALC_HEX24. The calcium ions shown in green stabilize the inter-triangle interfaces. **b-c**, Negative-stain TEM images showing salt-dependent reversible assembly between oligomers **(b)**, and arrays **(c)**. The inset in panel **b** shows a representative 2D-class average of the oligomer, and the inset in panel **c** shows the propagated array model. **d**, Negative-stain TEM 2D-class average of the designed array. **e**, Unstained HRTEM image of calcite nanocrystals templated by the oligomer. The shown lattice fringes with spacings of 3.02 and 2.41Å and interfringe angle of ∼53°, consistent with the calculated angle between calcite (104) and (110) planes. **f**, Unstained HRTEM image of calcite nanocrystals templated by the arrays. The inset FFT pattern shows an inter-reflection angle of ∼30° between spots indexed to the calcite (113) and (202) reflections. **g**, AFM image of mineralized arrays. **h**, High-magnification AFM image of the crystals templated on the protein arrays with angles consistent with the 102° angle of rhombohedral calcite. **i**, corresponding height profile measured along the red line in Panel g, showing step heights of approximately 4.5 nm.

Following NaCl-induced assembly, we transferred the arrays into pre-mineralization conditions by introducing 5 mM CaCl_2_ and removing NaCl through dialysis. Although stacking of the array sheets was observed during this process, the overall ordered array structure was preserved (**Fig. S6a**). In contrast, when protein was directly introduced into CaCl_2_-containing solution without NaCl-induced pre-assembly, no ordered arrays were observed and instead nonspecific aggregation occurred (**Fig. S6b)**.

CaCO_3_ mineralization of CALC_HEX24 was carried out in 5 mM CaCl_2_ and 5 mM NaHCO_3_ at a protein concentration of 0.014 mg/mL. We first examined mineralization by non-assembled CALC_HEX24. High-resolution transmission electron microscopy(HRTEM)revealed small nanoparticles approximately 5–10 nm in diameter (**Fig. 3e, Fig. S7a**). FFT analysis of the HRTEM region containing multiple nanoparticles showed spots corresponding to real-space d-spacings of 3.02 Å and 2.41 Å, indexed as the (104) and (110) reflections of the calcite-type phase, respectively (**Fig. S7b**). A representative high-resolution image further resolved lattice fringes with spacings of 3.02 Å and 2.41 Å, assigned to calcite (104) and (110) planes, with an interfringe angle of approximately 53° (**Fig. 3e**). We next examined mineralization by pre-assembled CALC_HEX24 arrays, and TEM imaging showed the formation of large crystals, up to ∼2 μm (**Fig. S8**). Further HRTEM and corresponding FFT analysis showed spots indexed as the calcite (113) and (202) reflections, with an inter-reflection angle of ∼30° in the FFT pattern. (**Fig. 3f**). By contrast, reactions lacking protein or containing negatively charged BSA produced vaterite-like morphologies (**Fig. S9**), consistent with previous reports^15^. AFM of the mineralized CALC_HEX24 shows that the protein arrays remain intact and exhibit layered structures on the substrate (**Fig. 3g**). Rhombohedral crystals with edge angles of approximately 102° were observed on the protein array layers (**Fig. 3h**), whereas no comparable structures were detected on exposed silicon wafer regions (**Fig. S10**). The height profile measured along with the pink line in **Fig. 3g** reveals two discrete height steps of 4.5 nm, consistent with stacking of two individual array layers (**Fig. 3i**). Together with the HRTEM analyses, these results show that CALC_HEX24 arrays induce localized, protein-templated calcite mineralization.

### Designed templating of carbonate mineral not found biological systems

We next sought to test whether our lattice-matching strategy could be extended beyond calcium carbonate to compounds not found in biological systems. We targeted CoCO_3_, as cobalt is a critical metal for energy-relevant materials, and protein-directed cobalt enrichment could provide a genetically encoded route to control cobalt-based composites with potential applications in electrocatalysis and photoelectrochemical energy conversion^11^. CoCO_3_ adopts the calcite-group rhombohedral/trigonal lattice, preserving the general carbonate sublattice organization found in calcite, but substitution of Ca^2+^ by the smaller Co^2+^ ion contracts the lattice, altering the spacing of surface-exposed interaction sites while maintaining an analogous carbonate arrangement.

To target the {101}^25^ facet of CoCO_3_ (**Fig. 4a**), we adapted the lattice-matched strategy used for ARAG64, positioning acidic side-chain carboxylates so that their terminal oxygen atoms matched carbonate oxygen positions on the mineral surface. Tip-atom-guided RFD2-RP was then used to scaffold these carboxylates at the ends of aspartate or glutamate residues, and designs capable of propagating along the {101} lattice without steric clashes were linked through a second diffusion step to create an extended protein-mineral interface. This workflow yielded COCA46, a designed repeat-protein scaffold for the CoCO_3_ {101} facet (**Fig. 4b**). COCA46 runs as a monomer in size-exclusion chromatography (SEC) (**Fig. S11**) with CD spectrum exhibiting alpha-helical folds at 25°C (**Fig. 4c**). During concentration, the pink color from cobalt ions remained with COCA46 in the retentate, suggesting ion binding (**Fig. S12**).

**Figure 4.**
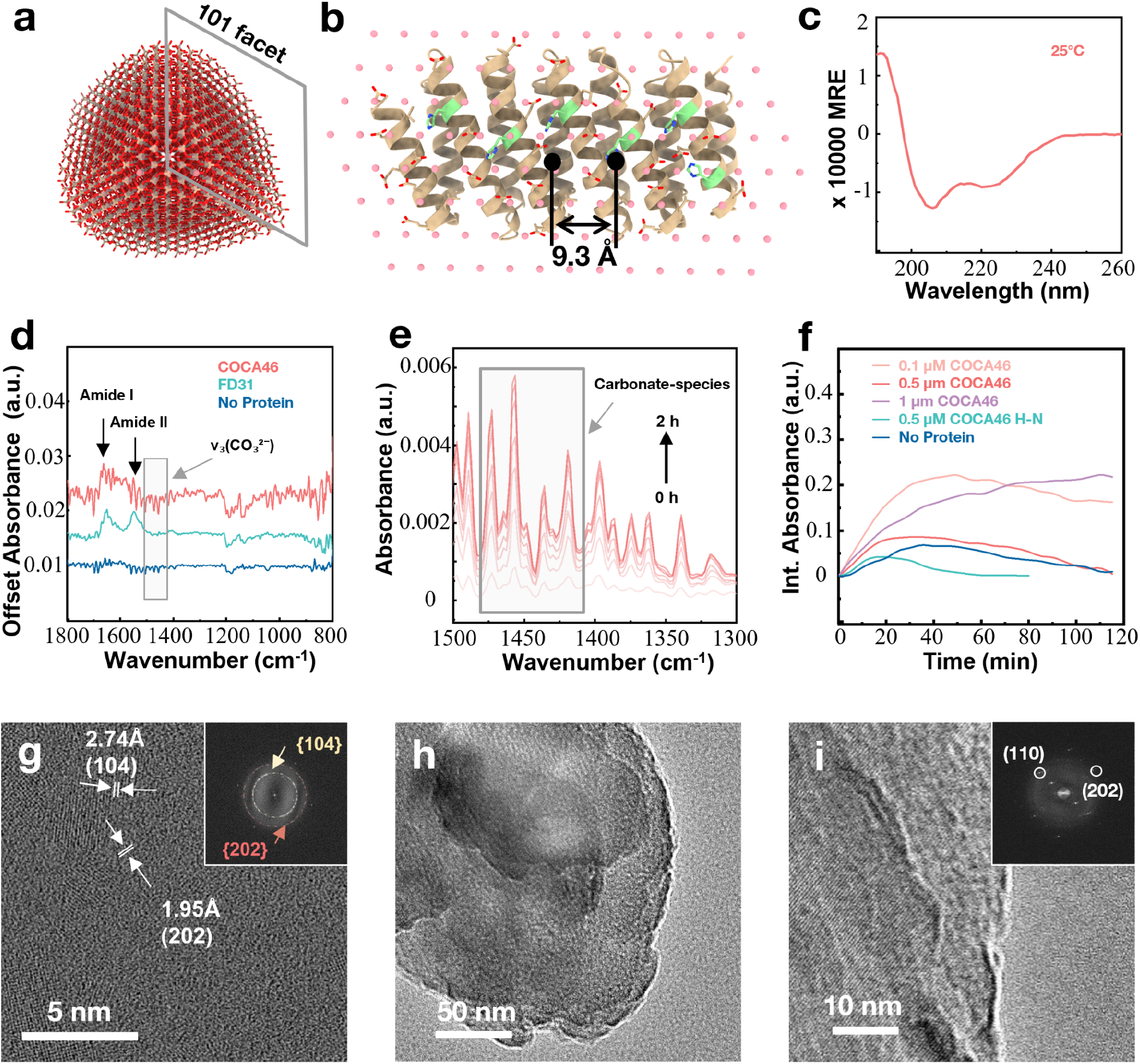
CoCO_3_ templated by COCA46 protein. **a**, Crystal structure of CoCO_3_, with an exposed {101} facet marked in grey. **b**, Design model of COCA46 protein targeting the CoCO_3_ {101} facet with designed binding motifs at a 9.3 Å lattice-matching spacing. **c**, CD spectrum of COCA46 protein shows α-helical folds. **d**, ATR-FTIR spectra collected after 2 hours of mineralization with 0.5 µM COCA46 (red), 0.5 µM FD31 (green), and the no-protein control (blue) in 1 mM CoCl_2_ and 1 mM NaHCO_3_ at pH 8.2 and 25°C. Protein-containing reactions show amide I and amide II bands at 1650 and 1540 cm^−1^, whereas these bands are absent from the no-protein control. The carbonate-associated vibrational region used for analysis is shaded. **e**, Representative time-resolved ATR-FTIR spectra showing the evolution of carbonate-associated peaks during mineralization from 0-2 hours. The shaded region (1410 to 1480 cm^−1^) indicates the carbonate ?_3_ integration window used for carbonate quantification plotted in panel **f. f**, Integrated (Int.) carbonate-associated signal as a function of time for CoCO_3_ mineralization with COCA46 protein at different concentrations (0.1, 0.5, and 1 µM), the 0.5 µM COCA46 H→N interface mutant, and the no-protein control. **g**, HRTEM image of CoCO3 nanocrystal after 3 hours of mineralization with 0.5 µM COCA46 in 1 mM CoCl_2_ and 1 mM NaHCO_3_ at pH 8.2 and 25°C. The lattice fringes with spacings of 2.74 and 1.95 Å are assigned to the CoCO_3_ (104) and (202) planes. The inset FFT shows the rings indexed to {104} and {202} reflections. **h-i**, TEM images of CoCO_3_ particles after 3 days of mineralization in the mineralization conditions of Panel g. The inset FFT panel **i** shows spots indexed as the CoCO_3_ (110) and (202) reflections.

COCA46 was incubated at 25°C with 1 mM CoCl_2_ and 1 mM NaHCO_3_, and time-resolved ATR-FTIR was used to monitor carbonate-associated signal evolution during mineralization (**Fig. 4d–f**). Protein-containing reactions showed clear amide I and amide II bands^34^, confirming the presence of protein during measurement, whereas these bands were absent from the no-protein control (**Fig. 4d**). The signal in the carbonate-associated vibrational regions^35,36^ was greater in the 0.5µM COCA46 case compared to 0.5 µM FD31 and no protein controls under the same precursor conditions (**Fig. 4d**). Quantification of integrated carbonate-associated signal profiles (**Fig. 4e**) revealed the concentration-dependent carbonate-species accumulation across 0.1, 0.5, or 1.0 µM COCA46 conditions, with 0.5 µM COCA46 H→N knockout mutant (**Table S1**), and no-protein condition (**Fig. 4f**) serving as controls. The 0.1 µM and 0.5 µM COCA46 conditions showed similar kinetic profiles-a rapid early accumulation followed by a gradual decrease in carbonate-associated signal, with the 0.5 µM condition producing a larger signal amplitude than the 0.1 µM condition. The 1.0 µM condition showed a more sustained carbonate-associated signal over the measurement period. Excess Co^2+^-binding sites could reduce free Co^2+^ availability and lower the effective supersaturation, thereby stabilizing carbonate-containing species before their conversion into larger mineral products.

Following ATR-FTIR measurements, we examined the resulting mineralization products by SEM and TEM. After 3 hours of mineralization, HRTEM of the 0.5 µM COCA46 condition revealed particles approximately 5 nm in diameter with lattice spacings of 2.74 Å and 1.95 Å, assigned to (104) plane and (202) reflection of calcite-structured CoCO_3_, respectively, with corresponding FFT rings indexed to the {104} and {202} reflections (**Fig. 4g, Fig. S13a**). Because the (202) reflection is a second-order associated with (101) plane, the appearance of (202) lattice fringes in HRTEM supports the engagement of the intended non-natural facet. Additional 2.74 Å fringes, intersected at ∼85°, are consistent with symmetry-related {104} planes (**Fig. S13b**), further supporting the assignment to calcite-structured CoCO_3_. Over 3 days of incubation, these particles grew into larger crystals containing cobalt (**Fig. S14a-b**). HRTEM image and FFT analysis of the 3-day products showed spots consistent with the (202) and (110) reflections of CoCO_3_ (**Fig. 4h-i**), indicating progressive structural ordering of the crystal phase. In contrast, the no-protein control (**Fig. S15a-b**) and the previously reported calcite-binding repeat protein FD31^15^ both yielded amorphous precipitates (**Fig. S16-17**), highlighting the role of the designed protein interface on directing CoCO_3_ mineralization. Disruption of the designed interface through the COCA46 H→N mutations led to amorphous or cobalt hydroxide particles after 3 days (**Fig. S18**), indicating the loss of CoCO_3_ templating capability. To further assess whether the mineralization capability is design-specific, we examined design COCA54 which targets the same {101} facet along a different spacing direction (**Fig. S19a-b**). COCA54 contains fewer histidines at the protein–mineral interface and a greater number of negatively charged side chains, and produces predominantly amorphous precipitates (**Fig. S20-21**). Taken together, these results indicate that productive mineral ordering depends on the precise side-chain chemistry presented at the protein–mineral interface.

### Design and characterization of CoCO_3_ templating dihedral cage

Having established that the lattice-matched protein interfaces can direct CoCO_3_ nucleation and growth, we next evaluated whether this design strategy could be extended to template and encapsulate CoCO_3_ in a defined crystal geometry. To address this, we designed a D3-symmetric protein architecture with interior surfaces matching the CoCO_3_ {110} facet lattice with LigandMPNN^26^. (**Fig. 5a**). Out of the 12 dihedral designs screened (**Table S1**), one, COCA_BRL11, was confirmed with negative-stain TEM to form the intended protein cages (**Fig. S22**). We succeeded in solving the crystal structure of COCA_BRL11, and found it to be in close agreement with the computational design model (1.99 Å RMSD) (**Fig. 5b; Table S2**) with a barrel-like assembly surrounding a hollow interior cavity.

**Figure 5.**
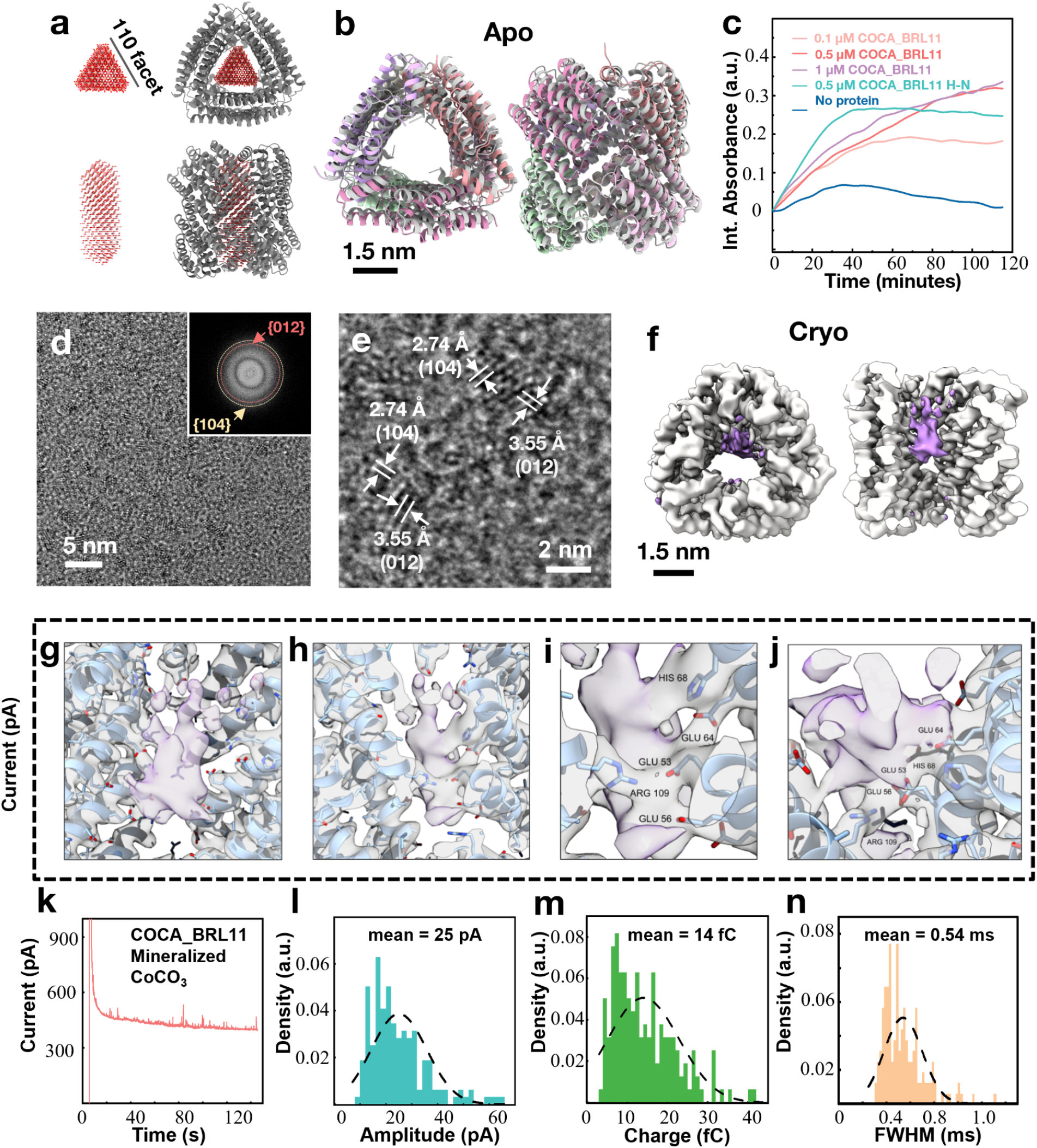
Cobalt carbonate templating and confinement by a designed protein cage. **a**, Design model of COCA_BRL11 protein, a D3-symmetric oligomer with protein interfaces that have lattices matched with the CoCO_3_ {110} facets. **b**, Apo crystal structure of COCA_BRL11 aligned to the design model (gray). **c**, Time-resolved FTIR as a function of time for CoCO_3_ mineralization with COCA_BRL11 protein at different concentrations (0.1, 0.5, and 1 µM), the 0.5 µM COCA_BRL11 H→N interface mutant, and the no-protein control in 1 mM CoCl_2_ and 1 mM NaHCO_3_ at 25°C and pH 8.2. **d**, Unstained HRTEM image of CoCO3 nanocrystals formed with COCA_BRL11 after 1 day mineralization. The inset FFT shows rings indexed to the {104} and {012} reflections. **e**, HRTEM image showing lattice fringes with spacings of 2.74 and 3.55 Å, assigned to CoCO_3_ (104) and (012) planes, respectively. **f**, Cryo-EM maps in top and cutaway-side views with a 3.46 Å resolution refined without symmetry (C1). CoCO_3_ and protein densities are shown in purple and gray, respectively. shown. **g**, Transparent cutaway-side view zoomed in Panel f. **h**, 180° view of Panel g showing the amino acid residues-mineral interfaces. **i**, Zoom in view of Panel h showing the labelled residues with continuous density with CoCO_3_. **j**, Top view of Panel i showing the amino acid residues-mineral interface 90° view of (i). **k**, Representative current–time trace of COCA_BRL11-mineralized CoCO_3_ particles recorded in 0.1 M KOH at 1.0 V versus Ag/AgCl using a carbon fiber microelectrode. **l–n**, Histograms of baseline-corrected peak amplitude (**l**), integrated charge (**m**), and FWHM (**n**) for isolated OER-associated current transients while excluding any signals attributed to the simultaneous collision of multiple particles or particle aggregates. Dashed curves are guides to the distributions. Values indicate mean values from the analyzed peaks.

Time-resolved ATR-FTIR measurements of COCA_BRL11 and the COCA_BRL11 H→N mutant under the mineralization conditions established for COCA46 (1 mM CoCl_2_ and 1 mM NaHCO_3_) showed distinct carbonate-associated kinetic profiles (**Fig. 5c, Fig. S23**). The mutant initially showed higher intensity, but was overtaken by original COCA_BRL11 design at equimolar protein concentrations after 80 minutes, suggesting that both cage-scaffolds enrich carbonate-species under the same conditions, but the designed histidine-mediated interface changes their subsequent evolution. We next examined the mineral products formed in the 0.5 µM COCA_BRL11 condition. After 3 hours, well-dispersed nanoparticles were observed (**Fig. S24a**), and HRTEM revealed lattice spacings consistent with the (110), (104), and (012) planes of CoCO_3_, confirming the growth of crystalline CoCO_3_ nanocrystals (**Fig. S24b**). After 3 days, these nanoparticles did not evolve into large single-crystals, but formed polycrystalline aggregates composed of small particles which SEM-EDX showed contained cobalt (**Fig. S25**). HRTEM imaging of these particles and FFT analysis revealed real-space lattice spacings and reflection features consistent with the (104), (113), (110), and (202) spacings of CoCO_3_ crystals (**Fig. S26**). In contrast, the COCA_BRL11 H→N mutant failed to produce CoCO_3_ crystals under the same conditions and instead yielded amorphous precipitates (**Fig. S27a-b**). Together, these results indicate that the designed histidine-mediated protein–mineral interface is required to direct crystalline CoCO_3_ formation.

To evaluate the potency of COCA_BRL11 in nucleating CoCO_3_, we experimented with lower supersaturation conditions with 0.25 mM CoCl_2_ and 0.25 mM NaHCO_3_. Under these conditions, 0.3 µM COCA_BRL11 promoted the formation of small crystalline particles within 1 day, with HRTEM and corresponding FFT revealed real-space lattice spacings and reflection rings assigned to CoCO_3_ (012) and (104) spacings (**Fig. 5d-e, Fig. S28**). In contrast, under the identical conditions, the no-protein control (**Fig. S27c**) and COCA46 (**Fig. S27d**) produced amorphous precipitates. The designed protein cage likely creates a spatially confined environment that locally enriches the ions and promotes CoCO3 nucleation.

To probe Co^2+^ binding within the designed scaffold, we first use cryo-TEM to determine a holo crystal structure of COCA_BRL11 after soaking them with CoCl2. The structure showed Co2+ incorporation predominantly at the intended mineral-facing interfaces of COCA_BRL11 (**Fig. S29a**), with the protein backbone closely matching the design model (2.32 Å RMSD) (**Fig. S29b**). These bound Co2+ suggest that COCA_BRL11 interfaces could direct the formation of cobalt-based inorganic phases by templating and enriching Co2+. We then prepared a monodisperse CoCO_3_-mineralized COCA_BRL11 population by size-exclusion chromatography (**Fig. S29c**) and analyzed it using cryo-EM. Representative micrographs and 2D class averages showed nanoparticles with a range of internal densities (**Fig. S29d**). The 3D reconstruction (**Fig. 5f**) had a 1.76 Å RMSD with the design model (**Fig. S29e**), supporting the retention of the overall D3 cage geometry after mineralization (**Fig. S30; Fig. S31; Table S3**), and shows additional density that cannot be attributed to the protein in its internal cavity–presumably from the CoCO_3_ nanocrystals–adjacent to the designed CoCO_3_ templating interfaces (**Fig. 5g-j**). These data support a model in which COCA_BRL11 acts as a protein container that templates and confines CoCO_3_ crystal formation within a defined oligomeric framework.

### Dihedral cage-templated CoCO_3_ forms electrocatalytically active protein–mineral composites

To evaluate catalytic function of the protein-cage-CoCO_3_ hybrid, we incubated 0.3 µM COCA_BRL11, 0.25 mM CoCl_2_ and 0.25 mM NaHCO_3_ in water for 1 day at 25°C to generate COCA_BRL11-mineralized CoCO_3_ particles for electrochemical testing. Electrochemical measurements showed that COCA_BRL11-mineralized CoCO_3_ sustained oxygen evolution reaction (OER) activity under alkaline conditions (**Fig. 5k**). The OER response varied with the concentration of COCA_BRL11-mineralized CoCO_3_ (**Fig. S32**), consistent with a particle-concentration-dependent contribution to the observed collision events. COCA_BRL11 alone showed no measurable activity above background, and CoCO_3_ formed without protein showed only low activity (**Fig. S33**). Several controls with the same mineralization conditions, including BSA, the COCA_BRL11 H→N mutant, and DHR46 showed initial OER responses but failed to sustain activity over time (**Fig. S33**). Event-level analysis gave mean peak amplitudes of 25 pA, integrated charges of 14 fC, and full width at half maximum (FWHM) values of 0.54 ms, indicating finite, sub-millisecond OER-associated collision events (**Fig. 5l–n, Fig S34**). These results suggest that the homogenous CoCO3 nanocrystals confined by COCA_BRL11 cages prolonged OER activity.

## Discussion

This work demonstrates that *de novo* protein design can control mineralization across multiple length scales and modes of organization, including polymorph selection, unbounded growth on reconfigurable protein arrays, and bounded geometric confinement within protein cages. ARAG64 selectively promotes aragonite while FD31 favors calcite under identical conditions that otherwise produced mixed phases, showing that precise chemical presentation on an ordered interface is sufficient to bias phase outcomes. COCA_BRL11 confined CoCO_3_ formation within the cage interior, producing uniform crystalline nanoparticles with the protein architecture remaining intact after mineralization. This result shows that designed protein structures can actively constrain and shape mineral growth rather than merely templating nucleation along a surface.

Previous efforts to template non-biological minerals such as ZnO and FeO have largely relied on threading metal-interacting residues onto flat repeat-protein scaffolds, followed by experimental screening or yeast-display selection against naturally occurring crystal faces^15,18^. More recently, lattice-matching strategies using oligomeric assemblies have been explored^17^, but these approaches produced crystals with heterogeneous sizes. Here, we move beyond these strategies by using RFD-based repeat-protein design to directly embed geometry-optimized atomic motifs into multi-helix scaffolds, enabling programmable targeting of metastable polymorphs and non-natural crystal facets without requiring high-throughput screening. This design logic extends from calcite and aragonite to CoCO_3_, a mineral with no known biological precedent, substantially broadening the scope of protein-directed mineralization. Moreover, confinement within the COCA_BRL11 cage yielded CoCO_3_ nanoparticles with remarkably coherent sizes. Together with previous demonstrations of mineral confinement in designed protein scaffolds, including hydroxyapatite and zinc oxide^17,18^, these results expand the possibility of using de novo protein architectures to template and confine the formation of coherent inorganic nanoparticles.

Overall, our results show that the polymorph selectivity and spatial control achieved by native mineralizing matrices can be reproduced and extended using ordered, programmable protein scaffolds designed from scratch. Thus the results establish de novo design as a tractable route to recreate some of the most distinctive features of biological mineralization. Beyond fundamental insight, the framework carries practical significance: genetically encodable control over carbonate mineralization has potential applications in biocement and carbon management^37^, while the electrocatalytic activity of COCA_BRL11-derived cobalt carbonate particles demonstrates that designed biomineralization scaffolds can produce functional hybrid bioinorganic materials.

## Acknowledgments

PSK thanks C. Fries, N. Hanikel, A. Favor, S. Wang, S. Rankovic, and Z. Jones for discussion on 2d arrays. PSK thanks A. Favor and A. Kubaney for discussion on RFdiffusion and MPNN. PSK thanks P. Lund-Anderson for sharing his 10 mM Co(II)Cl_2_ stock. PSK thanks J. Decarreau for guidance on the IN Cell Analyzer. PSK thanks A. Swartz, E. Huddy, Y. Politanska and S. Moroz and the rest of Baker lab members for general discussions. We thank the IPD lab management team led by K. VanWormer, IPD IT team led by L. Goldschmidt, IPD Electron Microscopy Research Core (EMRC) led by A. Borst for technical support. Joel Quispe, Fangfang Zhang, and Wentao Jiang for management of the University of

Washington Cryo-EM facilities. We thank Howard Hughes Medical Institute (DB). Design, production, and characterization of the aragonite and CoCO_3_ targeting designs was performed at the IPD in the Biochemistry Department at UW through support from the DOE SC BES, as part of the Energy Frontier Research Centers (EFRC): CSSAS, The Center for the Science of Synthesis Across Scales, under Award DE-SC0019288. AFM, ATR-FTIR, SEM, EDS/EDX, HRTEM characterizations were also supported by CSSAS at PNNL through FWP 72448. PNNL is a multiprogram national laboratory operated for DOE by Battelle under Contract No. DE-AC05-76RL01830.

This research used resources 17-ID-2 (FMX) of the National Synchrotron Light Source II, a U.S. Department of Energy (DOE) Office of Science User Facility operated for the DOE Office of Science by Brookhaven National Laboratory under Contract No. DE-SC0012704. The Center for BioMolecular Structure (CBMS) is primarily supported by the National Institutes of Health, National Institute of General Medical Sciences (NIGMS) through a Center Core P30 Grant (P30GM133893), and by the DOE Office of Biological and Environmental Research (KP1605010).

## Funding

Howard Hughes Medical Institute

The Audacious Project at the Institute for Protein Design

The Open Philanthropy Project Improving Protein Design Fund

This material is based on work supported by the Air Force Office of Scientific Research under award number FA9550-22-1-0506

This material was based upon work supported by the U.S. Department of Energy, Office of Science, Office of Basic Energy Sciences, as part of the Energy Frontier Research Centers program: The Center for the Science of Synthesis Across Scales under Award Number DE-SC0019288.

## Author contributions

Conceptualization: PSK, XL, LTY, HP, JDY, DB

Methodology: PSK, XL, LTY, HP, CW, AJB, CCB, THL, AK, HN, JM, KDC, BC, YH, ZM,DL

Investigation: PSK, XL, LTY, HP, CW, AJB, CCB, THL, AK, HN, JM, KDC

Data processing: PSK, XL, LTY, CW, JM, HN, AK, AJB, SZ

Visualization: PSK, XL, LTY, CW, JM, HN, AK, AJB, SZ

Funding acquisition: SZ, JDY, BZ, BMC, DB Supervision: BZ, BMC, SZ, AKB, JDY, DB

Writing – original draft: PSK, XL, LTY, HP, CW, DB

Writing – review & editing: All authors

Competing interests: Authors declare that they have no competing interests. Data and materials availability:

All data are available in the main text or the supplementary materials. Scripts for RFdiffusion 2 are available at https://github.com/RosettaCommons/RFdiffusion2. Scripts for RFdiffusion for symmetric motif scaffolding are available at https://github.com/RosettaCommons/RFdiffusion. Scripts for ProteinMPNN and LigandMPNN are available at https://github.com/dauparas/LigandMPNN. Scripts for AlphaFold2 and AlphaFold3 are available at https://github.com/google-deepmind/alphafold and https://github.com/google-deepmind/alphafold3 respectively. All detailed scripts for design pipelines and fine-tuned weights will be available on github at the time of publication.

## References

1. Suzuki, M. et al. An Acidic Matrix Protein, Pif, Is a Key Macromolecule for Nacre Formation. Science 325, 1388–1390 (2009).

2. Nudelman, F. et al. The role of collagen in bone apatite formation in the presence of hydroxyapatite nucleation inhibitors. Nat. Mater. 9, 1004–1009 (2010).

3. Chen, C.-L., Bromley, K. M., Moradian-Oldak, J. & DeYoreo, J. J. In situ AFM Study of Amelogenin Assembly and Disassembly Dynamics on Charged Surfaces Provides Insights on Matrix Protein Self-Assembly. J. Am. Chem. Soc. 133, 17406–17413 (2011).

4. Siponen, M. I. et al. Structural insight into magnetochrome-mediated magnetite biomineralization. Nature 502, 681–684 (2013).

5. Bahn, S. Y., Jo, B. H., Choi, Y. S. & Cha, H. J. Control of nacre biomineralization by Pif80 in pearl oyster. Sci. Adv. 3, e1700765 (2017).

6. Sun, J. & Bhushan, B. Hierarchical structure and mechanical properties of nacre: a review. RSC Adv. 2, 7617 (2012).

7. Falini, G., Albeck, S., Weiner, S. & Addadi, L. Control of Aragonite or Calcite Polymorphism by Mollusk Shell Macromolecules. Science 271, 67–69 (1996).

8. Liang, H., Liu, Y., Tian, B., Li, Z. & Ou, H. A sustainable production of biocement via microbially induced calcium carbonate precipitation. Int. Biodeterior. Biodegrad. 172, 105422 (2022).

9. Magnetically sensitive proteins could lead to new imaging tools and remote-controlled drugs. 10.1126/science.zzfau8s (2026).

10. Tang, S., Dong, Z., Ke, X., Luo, J. & Li, J. Advances in biomineralization-inspired materials for hard tissue repair. Int. J. Oral Sci. 13, 42 (2021).

11. Zhang, X., Yi, H., An, Q. & Song, S. Recent advances in engineering cobalt carbonate hydroxide for enhanced alkaline water splitting. J. Alloys Compd. 887, 161405 (2021).

12. Zhang, S., Pyles, H., Baker, D. & De Yoreo, J. J. Designing protein–material interfaces. MRS Bull. 50, 1454–1464 (2025).

13. Boskey, A. L. & Villarreal-Ramirez, E. Intrinsically disordered proteins and biomineralization. Matrix Biol. 52–54, 43–59 (2016).

14. Yang, W., Wang, S. & Lee, G. R. The past, present and future of de novo protein design. Nature 652, 1139–1152 (2026).

15. Davila-Hernandez, F. A. et al. Directing polymorph specific calcium carbonate formation with de novo protein templates. Nat. Commun. 14, 8191 (2023).

16. Voets, I. K. From ice-binding proteins to bio-inspired antifreeze materials. Soft Matter 13, 4808–4823 (2017).

17. Yu, L. T. et al. Templating and confining calcium phosphate mineralization within designed protein assemblies. Preprint at 10.64898/2026.01.14.699524 (2026).

18. Saragovi, A. et al. De novo design of metal-oxide templating proteins. Preprint at 10.1101/2024.06.24.600095 (2024).

19. De Haas, R. J. et al. Inhibition of ice recrystallization with designed twistless helical repeat proteins. Proc. Natl. Acad. Sci. 122, e2514871122 (2025).

20. Ridgwell, A. & Zeebe, R. The role of the global carbonate cycle in the regulation and evolution of the Earth system. Earth Planet. Sci. Lett. 234, 299–315 (2005).

21. McDonald, K., Khajuria, D. K. & Li, B. Potential healthcare applications of metal carbonate nanoparticles produced from emerging carbon capture technologies. Nanomed. 21, 921–923 (2026).

22. Ahern, W. et al. Atom-level enzyme active site scaffolding using RFdiffusion2. Nat. Methods https://doi.org/10.1038/s41592-025-02975-x (2025) doi:10.1038/s41592-025-02975-x.

23. Berman, H. M. The Protein Data Bank. Nucleic Acids Res. 28, 235–242 (2000).

24. Varadi, M. et al. AlphaFold Protein Structure Database: massively expanding the structural coverage of protein-sequence space with high-accuracy models. Nucleic Acids Res. 50, D439–D444 (2022).

25. Caspi, E. N., Pokroy, B., Lee, P. L., Quintana, J. P. & Zolotoyabko, E. On the structure of aragonite. Acta Crystallogr. B 61, 129–132 (2005).

26. Dauparas, J. et al. Atomic context-conditioned protein sequence design using LigandMPNN. Nat. Methods 22, 717–723 (2025).

27. Jumper, J. et al. Highly accurate protein structure prediction with AlphaFold. Nature 596, 583–589 (2021).

28. Jiang, B., Jain, A., Lu, Y. & Hoag, S. W. Probing Thermal Stability of Proteins with Temperature Scanning Viscometer. Mol. Pharm. 16, 3687–3693 (2019).

29. Tran, L. et al. Design of Orthogonal Far-Red, Orange and Green Fluorophore-binding Proteins for Multiplex Imaging. Preprint at 10.1101/2025.08.03.668343 (2025).

30. Dauparas, J. et al. Robust deep learning–based protein sequence design using ProteinMPNN. Science 378, 49–56 (2022).

31. Punjani, A., Rubinstein, J. L., Fleet, D. J. & Brubaker, M. A. cryoSPARC: algorithms for rapid unsupervised cryo-EM structure determination. Nat. Methods 14, 290–296 (2017).

32. Gonen, S., DiMaio, F., Gonen, T. & Baker, D. Design of ordered two-dimensional arrays mediated by noncovalent protein-protein interfaces. Science 348, 1365–1368 (2015).

33. Ben-Sasson, A. J. et al. Design of biologically active binary protein 2D materials. Nature 589, 468–473 (2021).

34. Sizeland, K. H. et al. Nanostructure of electrospun collagen: Do electrospun collagen fibers form native structures? Materialia 3, 90–96 (2018).

35. Henry, D. G., Watson, J. S. & John, C. M. Assessing and calibrating the ATR-FTIR approach as a carbonate rock characterization tool. Sediment. Geol. 347, 36–52 (2017).

36. Grunenwald, A. et al. Revisiting carbonate quantification in apatite (bio)minerals: a validated FTIR methodology. J. Archaeol. Sci. 49, 134–141 (2014).

37. Beatty, D. N., Williams, S. L. & Srubar, W. V. Biomineralized Materials for Sustainable and Durable Construction. Annu. Rev. Mater. Res. 52, 411–439 (2022).

38. Steinegger, M. & Söding, J. MMseqs2 enables sensitive protein sequence searching for the analysis of massive data sets. Nat. Biotechnol. 35, 1026–1028 (2017).

39. Gražulis, S. et al. Crystallography Open Database (COD): an open-access collection of crystal structures and platform for world-wide collaboration. Nucleic Acids Res. 40, D420–D427 (2012).

40. Abramson, J. et al. Accurate structure prediction of biomolecular interactions with AlphaFold 3. Nature 630, 493–500 (2024).

41. Dang, B. et al. SNAC-tag for sequence-specific chemical protein cleavage. Nat. Methods 16, 319–322 (2019).

42. Studier, F. W. Protein production by auto-induction in high-density shaking cultures. Protein Expr. Purif. 41, 207–234 (2005).

43. Nannenga, B. L., Iadanza, M. G., Vollmar, B. S. & Gonen, T. Overview of Electron Crystallography of Membrane Proteins: Crystallization and Screening Strategies Using Negative Stain Electron Microscopy. Curr. Protoc. Protein Sci. 72, (2013).

44. Mastronarde, D. N. SerialEM: A Program for Automated Tilt Series Acquisition on Tecnai Microscopes Using Prediction of Specimen Position. Microsc. Microanal. 9, 1182–1183 (2003).

45. Sun, M. et al. Practical considerations for using K3 cameras in CDS mode for high-resolution and high-throughput single particle cryo-EM. J. Struct. Biol. 213, 107745 (2021).

46. Punjani, A., Rubinstein, J. L., Fleet, D. J. & Brubaker, M. A. cryoSPARC: algorithms for rapid unsupervised cryo-EM structure determination. Nat. Methods 14, 290–296 (2017).

47. Kim, K., Li, H. & Clarke, O. B. High-resolution ab initio reconstruction enables cryo-EM structure determination of small particles. Preprint at 10.1101/2025.09.08.674935 (2025).

48. Pettersen, E. F. et al. UCSF CHIMERAX : Structure visualization for researchers, educators, and developers. Protein Sci. 30, 70–82 (2021).

49. Emsley, P. & Cowtan, K. Coot : model-building tools for molecular graphics. Acta Crystallogr. D Biol. Crystallogr. 60, 2126–2132 (2004).

50. Emsley, P., Lohkamp, B., Scott, W. G. & Cowtan, K. Features and development of Coot. Acta Crystallogr. D Biol. Crystallogr. 66, 486–501 (2010).

51. Brown, A. et al. Tools for macromolecular model building and refinement into electron cryo-microscopy reconstructions. Acta Crystallogr. D Biol. Crystallogr. 71, 136–153 (2015).

52. Croll, T. I. ISOLDE : a physically realistic environment for model building into low-resolution electron-density maps. Acta Crystallogr. Sect. Struct. Biol. 74, 519–530 (2018).

53. Adams, P. D. et al. The Phenix software for automated determination of macromolecular structures. Methods 55, 94–106 (2011).

54. Williams, C. J. et al. MolProbity: More and better reference data for improved all-atom structure validation. Protein Sci. 27, 293–315 (2018).

55. Winter, G. xia2 : an expert system for macromolecular crystallography data reduction. J. Appl. Crystallogr. 43, 186–190 (2010).

56. Winter, G. et al. DIALS : implementation and evaluation of a new integration package. Acta Crystallogr. Sect. Struct. Biol. 74, 85–97 (2018).

57. Winn, M. D. et al. Overview of the CCP 4 suite and current developments. Acta Crystallogr. D Biol. Crystallogr. 67, 235–242 (2011).

58. Wojdyr, M., Keegan, R., Winter, G. & Ashton, A. DIMPLE - a pipeline for the rapid generation of difference maps from protein crystals with putatively bound ligands. Acta Crystallogr. A 69, s299–s299 (2013).

59. Murshudov, G. N. et al. REFMAC 5 for the refinement of macromolecular crystal structures. Acta Crystallogr. D Biol. Crystallogr. 67, 355–367 (2011).

60. Smart, O. S. et al. Exploiting structure similarity in refinement: automated NCS and target-structure restraints in BUSTER. Acta Crystallogr. D Biol. Crystallogr. 68, 368–380 (2012).

61. Joosten, R. P., Long, F., Murshudov, G. N. & Perrakis, A. The PDB_REDO server for macromolecular structure model optimization. IUCrJ 1, 213–220 (2014).

